# The majority of histone acetylation is a consequence of transcription

**DOI:** 10.1101/785998

**Authors:** Benjamin J.E. Martin, Julie Brind’Amour, Kristoffer N. Jensen, Anastasia Kuzmin, Zhen Cheng Liu, Matthew Lorincz, LeAnn J. Howe

**Author notes:** **Materials & Correspondence**: Correspondence and requests for materials should be addressed to.

## Abstract

Histone acetylation is a ubiquitous hallmark of transcriptional activity, but whether the link is of a causal or consequential nature is still a matter of debate. In this study we resolve this question. Using both immunoblot analysis and chromatin immunoprecipitation-sequencing (ChIP-seq) in *S. cerevisiae*, we show that the majority of histone acetylation is dependent on transcription. Loss of histone H4 acetylation upon transcription inhibition is partially explained by depletion of histone acetyltransferases (HATs) from gene bodies, implicating transcription in HAT targeting. Despite this, HAT occupancy alone poorly predicts histone acetylation, suggesting that HAT activity is regulated at a step post-recruitment. Collectively, these data show that the majority of histone acetylation is a consequence of RNAPII promoting both the recruitment and activity of histone acetyltransferases.

## Introduction

Lysine acetylation of histone amino terminal tails has been linked to gene expression for many decades^1^. More recently, genome-wide localization studies across eukaryotes, including yeast and mammals, revealed that histone tail acetylation primarily occurs at the promoters and 5’ ends of transcribed genes^2^. Although some forms of acetylation have been referred to as “global” and “non-targeted”^3^, genome-wide occupancy studies show that histone acetylation levels correlate strongly with transcription, implicating a causal relationship between the two.

Histone acetylation is catalyzed by conserved histone acetyltransferases (HATs), generally consisting of a catalytic subunit complexed with auxiliary proteins required for enzymatic activity and targeting^4^. Most HAT complexes have relatively low substrate specificity and modify multiple lysine residues within either H3 or H4. Thus, histone acetylation sites within H3 and H4 generally show similar distributions^2^, and mutations of histone lysine residues, with the exception of H4K16, result in comparable changes in gene expression^5^. Histone acetylation is a dynamic mark due to the activity of histone deacetylase complexes (HDACs). Similar to HATs, HDACs generally exist as multi-protein complexes with a catalytic subunit that can deacetylate multiple lysines on one or more histones^6^.

In *S. cerevisiae*, the most well characterized proteins with lysine acetyltransferase activity are Gcn5 and Esa1, which are the catalytic subunit of multiple HAT complexes, including the H3-specific HATs, SAGA and ADA for Gcn5, and the H4-specific HATs, NuA4 and Piccolo for Esa1^4^. SAGA and NuA4 are targeted to gene promoters via an interaction between a shared subunit, Tra1, and DNA-bound transcription activators^7–9^, which is thought to target acetylation of nucleosomes flanking promoters. This, together with the observation that Gcn5 and Esa1 are required for transcription of multiple genes^10–13^, has led to the widely accepted model that histone acetylation acts “upstream” of transcription initiation. It should be noted however that, in addition to canonical histones, HATs have been shown to acetylate many non-histone proteins involved in transcription initiation^14^. As such, whether SAGA and NuA4 activate transcription primarily through acetylation of the core histones remains uncertain.

While the current model is widely accepted, there are several examples of histone acetylation being deposited as a consequence of transcription. Numerous HATs interact with co-transcriptional H3K4me3^15–18^ and the phosphorylated carboxy-terminal domain (CTD) of RNAPII^12, 19^. Additionally, co-transcriptional histone exchange mediates incorporation of acetylated histones into nucleosomes within transcribed regions^20^, and recent work has shown that RNA can promote the activity of CBP at enhancers^21^. Taken together these observations suggest that histone acetylation can also be a consequence of the transcription process. However, the contribution of these pathways to histone acetylation patterns remains unknown.

In this study we sought to determine the relative contributions of the “causal” vs “consequential” pathways for targeting histone acetylation to transcribed genes. We first found that inhibition of transcription results in rapid histone deacetylation in both yeast and mouse embryonic stem cells (mESCs), demonstrating that histone acetylation is a consequence of transcription. Loss of RNAPII also results in redistribution of Epl1, a subunit of the NuA4 and Piccolo HATs, from coding regions to promoters, consistent with the transcription-independent targeting of HATs by activators and transcription-dependent targeting of HATs by RNAPII. However, despite the co-dependency of histone acetylation and HAT recruitment on RNAPII, we found that HAT recruitment alone is insufficient to mediate acetylation of the associated nucleosomes, indicative of post-recruitment regulation of acetyltransferase activity.

## Results

### The majority of histone acetylation is a consequence of transcription

Despite the well-known correlation between histone acetylation and transcription, whether this PTM is primarily a cause or consequence of transcription has not been definitively tested. We therefore sought to assess the dependence of histone acetylation on RNAPII by inhibiting transcription in *S. cerevisiae*. Previous studies have used the *rpb1-1* temperature sensitive mutant to disrupt transcription^22^. However more recent experiments have suggested that this mutant does not directly inhibit RNAPII, as shifting the mutant to the restrictive temperature has minimal effects on transcript synthesis^23^ and does not lead to rapid dissociation of RNAPII from gene bodies^24^. To achieve effective inhibition of RNAPII, we therefore treated cells with 1,10 phenanthroline monohydrate (1,10-pt), which has been shown to rapidly inhibit transcript synthesis^23^. Confirming efficient transcription inhibition by 1,10-pt, we observed a global loss of RNAPII serine 5 CTD phosphorylation by immunoblot analysis (Figure 1A) and rapid alterations in RNAPII distribution as determined by ChIP-seq (Figure 1B). Immunoblot analysis of yeast whole cell extracts showed that within 15 minutes of transcription inhibition, a broad range of H3 and H4 acetylation marks were rapidly lost (Figure 1A). Similar deacetylation was observed following, treatment with the transcription inhibitor thiolutin (Figure S1A), or degradation of Rpb2, the second largest subunit of RNAPII, using an auxin inducible degron (Figure S1B). Notably, loss of acetylation was dependent on the histone deacetylase Rpd3 (Figure S1C) and could be blocked by prior treatment with the HDAC inhibitor TSA (Figure S1D), confirming active deacetylation upon transcription inhibition. Histone acetylation loss was due to disruption of HAT activity, rather than increased HDAC activity, as treatment with TSA following 1,10-pt treatment failed to restore histone acetylation (Figure S1E). HATs are conserved throughout eukaryota and thus it is likely that acetylation is dependent on transcription in other organisms. Indeed, we found that inhibition of transcription by actinomycin D in mESCs resulted in global loss of H3K9ac and H3K27ac (Figure 1C). Thus, in yeast and in mouse cells, the majority of histone acetylation is dependent on transcription.

**Figure 1.**
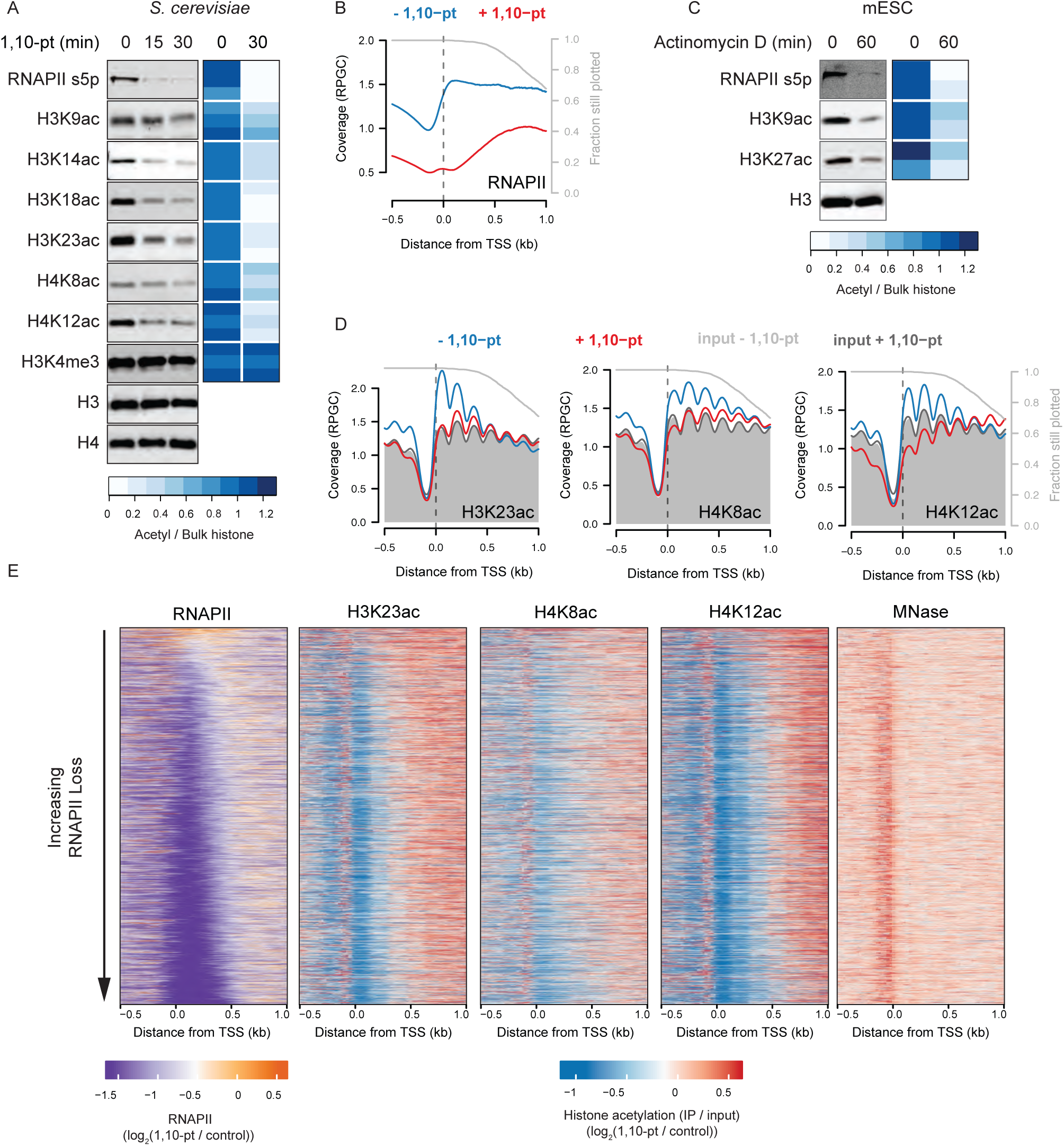
The majority of histone acetylation is dependent on transcription. **A**. Whole cell extracts from *S. cerevisiae* cells before and after treatment with 1,10-pt were subjected to immunoblot analysis with the indicated antibodies. Representative blots (left panel) and quantification of three independent replicates (right panel) are shown. For quantification, histone PTM signals were normalized to total histone H4 levels and expressed as a heatmap with the scale below. **B**. Average profile of RNAPII (Rpb3 ChIP-seq^27^) at 3862 transcribed genes (defined by RNAPII ChIP-seq binding) aligned by the TSS before and after a 15 minute treatment with 1,10-pt. Only data until the polyadenylation site (PAS) was included, and the grey line represents the fraction of genes still being plotted. RPGC, reads per genomic coverage; TSS, transcription start site. **C**. Nuclear extracts from mouse ESCs before and after treatment with actinomycin D were subjected to immunoblot analysis with the indicated antibodies. Representative blots (left panel) and quantification of two independent replicates (right panel) are shown. For quantification, histone acetylation signals were normalized to total histone H3 levels and expressed as a heatmap. **D**. Average profile of H3K23ac, H4K8ac, and H4K12ac (MNase ChIP-seq) at 3862 transcribed genes aligned by the TSS before and after a 15-minute treatment with 1,10-pt. Inputs from untreated (shaded grey) and 1,10-pt (dark grey line) are shown. **E**. Heatmaps representing the fold change (log_2_) following transcription inhibition for RNAPII (Rpb3 ChIP-seq), nucleosome occupancy (MNase-seq), and nucleosome-normalized H3K23ac, H4K8ac, and H4K12ac (MNase ChIP-seq) signal at all 5133 genes aligned by the TSS. Heatmaps are ordered by the change in RNAPII occupancy with the scale below.

To confirm that histone acetylation loss is a direct consequence of transcription inhibition, we treated *S. cerevisiae* cells with 1,10-pt for 15 minutes and performed ChIP-seq for H3K23ac, H4K8ac, and H4K12ac. Consistent with previous studies^25, 26^, we used non-transcribed regions to account for global changes in ChIP-seq experiments (see methods). While the mechanism of action for transcription inhibition by 1,10-pt has not been extensively characterized, Figure 1B and our previous work^27^ indicate inhibition of both transcription initiation and elongation. Histone acetylation was likewise lost following transcription inhibition (Figure 1D), while no major changes to nucleosome occupancy or positioning were observed following the short transcription inhibition performed here (Figures 1D and E). Importantly, heatmaps of the log_2_ fold change in ChIP-seq signal upon 1,10-pt treatment showed that loss of histone acetylation upon transcription inhibition mirrored that of RNAPII loss (Figure 1E). Additionally, histone deacetylation was limited to nucleosomes that lost RNAPII upon 1,10-pt treatment (Figure S1F). In contrast, regions with more stable RNAPII, including the 3’ ends of genes, showed slight increases in histone acetylation (Figures 1D and S1F). While this may result from enhanced HAT targeting by RNAPII displaced from other loci, we cannot rule out the possibility that our scaling approach did not fully account for global decreases in histone acetylation. Irrespective, we find that H3K23ac, H4K8ac, and H4K12ac were primarily deacetylated at regions that lost RNAPII upon transcription inhibition, indicative of a direct effect. Collectively these results demonstrate that the majority of histone acetylation is targeted as a consequence of transcription, which is inconsistent with the existing model that histone acetylation is targeted to active genes primarily via the interaction of HATs with transcription activators.

### Transcription promotes the interaction of H4-specific HATs with chromatin

The simplest explanation for the transcription dependence of histone acetylation is that RNAPII is required to target HATs to transcribed genes. To test this hypothesis we mapped Epl1, a common subunit of Esa1-dependent HATs^28^, by ChIP-seq prior to and following 15 minutes transcription inhibition with 1,10-pt. In the absence of drug, Epl1 was enriched over gene bodies, but depleted in the ∼200 bp upstream of the transcription start site (TSS) (Figure 2A). Following transcription inhibition Epl1 binding was reduced over the 5’ ends of gene bodies and increased upstream of the TSS (Figure 2A). The reduced Epl1 binding over gene bodies mirrored RNAPII loss (Figure 2B) indicating that RNAPII is required for targeting of HATs to transcribed chromatin.

**Figure 2.**
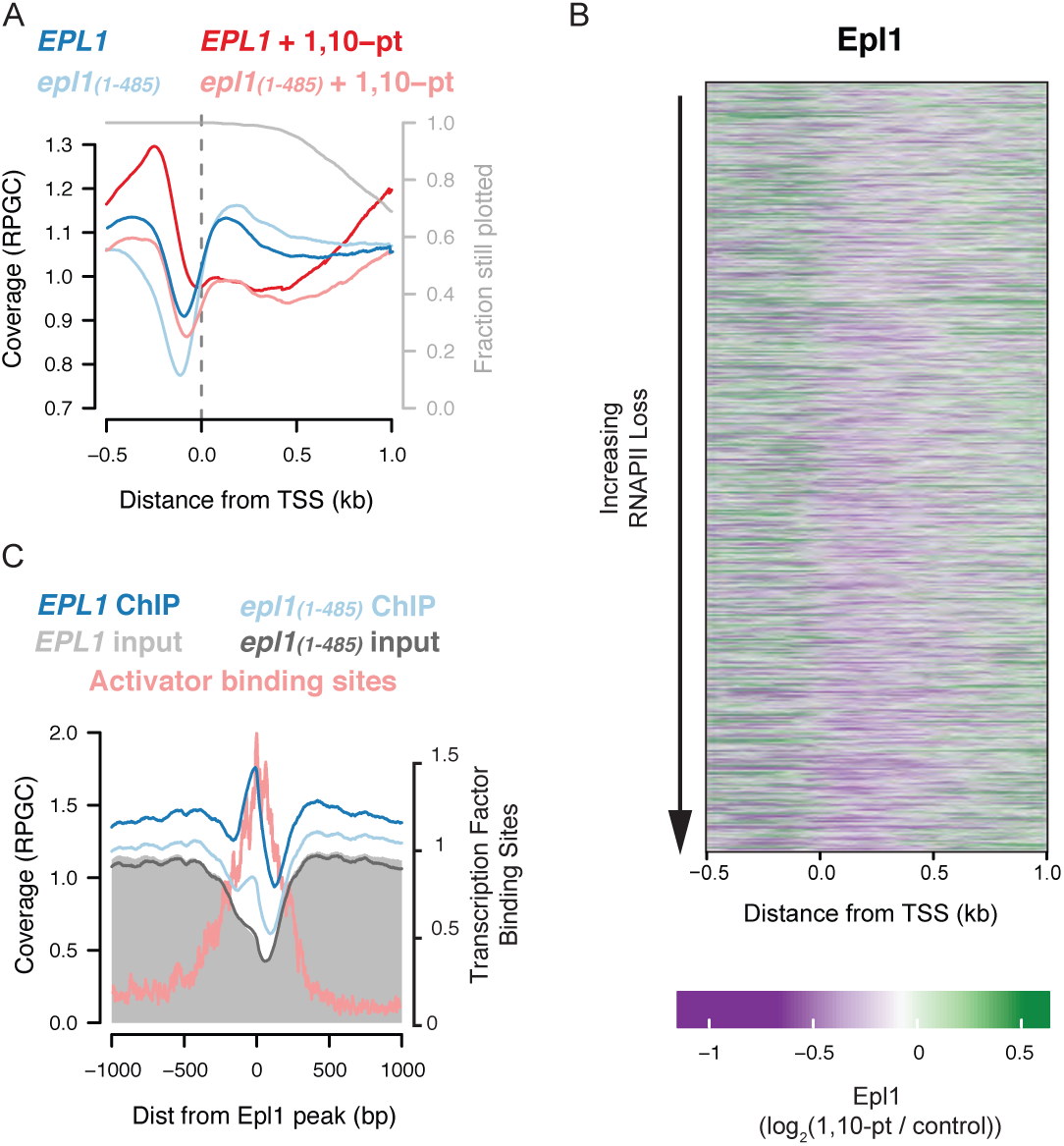
Transcription promotes the interaction of H4-specific HATs with chromatin. Average profile of Epl1 or Epl1_1-485_ (ChIP-seq from sonicated extracts) at 3520 transcribed genes aligned by the TSS before and after a 15-minute treatment with 1,10-pt. Only data until the PAS was included, and the grey line represents the fraction of genes still being plotted. RPGC, reads per genomic coverage; TSS, transcription start site. **B.** Heatmap representing the fold change (log_2_) following transcription inhibition for Epl1 ChIP-seq signal at 5133 genes aligned by the TSS. Heatmaps are ordered by the change in RNAPII occupancy, as in Figure 1E. **C**. Average profile of Epl1 and Epl1_1-485_ (ChIP-seq from MNase-treated extracts) relative to the center of 562 regions showing strong Epl1 peaks. Inputs from Epl1 (shaded grey) and Epl1_1-485_ (grey line) strains are shown.

Increased Epl1 binding upstream of the TSS upon transcription inhibition is consistent with recruitment of HATs displaced from gene bodies to promoters via interaction with transcription activators. To examine this more closely, we repeated the analysis on untreated cells using chromatin fragmented by micrococcal nuclease, which was previously shown to detect non-histone protein complexes bound to promoters^29^. This approach allowed is to identify 562 genes with strong peaks of Epl1 that were, on average, 200 bp upstream of the TSS (Figure 2C). Note that these peaks contained sub-nucleosome sized DNA fragments that did not precipitate with anti-acetyl-histone antibodies (Figures S2A and S2B), and thus were unlikely to represent nucleosomes. Moreover, these peaks overlapped regions with transcription factor binding sites (Figure 2C)^30^, indicative of Epl1 targeting by transcription activators. To further confirm that promoter-bound Epl1 was due to activator targeting, we repeated Elp1 ChIP-seq, from both sonicated and MNase-digested chromatin, in a strain expressing Epl1 lacking the 348 C-terminal amino acids (Epl1_1-485_). While this truncated protein still interacts with Esa1 and Yng2, it fails to co-purify with the remaining NuA4 subunits, including Tra1, which is required for the interaction of NuA4 with transcription activators^28^. As shown in Figures 2A (sonicated ChIP-seq) and 2C (MNase ChIP-seq), truncation of Epl1 reduces binding over promoters consistent with loss of activator targeting in this mutant. Notably, truncation did not decrease Epl1 binding to gene bodies suggesting that the activator interaction is not required for HAT binding in these regions (Figures 2A). Together these results indicate that Epl1-containing HATs are targeted to promoters and the bodies of transcribed genes, with the former likely through interaction with transcription activators.

RNAPII-dependent targeting of Esa1 to gene bodies has been proposed to occur through recognition of H3K4 methylation by the PHD finger of Yng2^31, 32^ and preferential interaction of NuA4 with the serine 5 phosphorylated form of RNAPII^12^. However, H3K4me3 was stable to transcription inhibition (Figure 1A), demonstrating that H3K4me3 alone cannot mediate Epl1 association with gene bodies. Furthermore, only subtle effects on bulk H4 acetylation resulted from deletion of the PHD finger of Yng2 (Figure S3A) or the gene encoding the sole H3K4 methyltransferase, Set1 (Figure S3B and C). Additionally, using an analog-sensitive *KIN28* mutant, we show that deposition of global acetylation was not dependent on RNAPII serine 5 phosphorylation by Kin28 (Figure S3D), either alone or in combination with loss of H3K4 methylation (Figure S3E), demonstrating that RNAPII-dependent recruitment of Epl1 to gene bodies is through a novel pathway. Although the exact mechanism remains to be identified, these results show that RNAPII targets histone acetylation, at least in part, through promoting the interaction of HATs with chromatin.

### The activity of histone acetyltransferases is regulated post-recruitment

The transcription-dependence of both histone acetylation and Elp1 occupancy suggests that the two should be linked, however close analysis reveals that HAT occupancy alone is insufficient to explain histone acetylation. As a first indication of this, the increased binding of Epl1 upstream of TSSs upon transcription inhibition (Figures 2A and C) was not associated with increased H4K8 or H4K12 acetylation (Figures 1D and S2B), suggesting that activator targeting of HATs is not sufficient for histone acetylation *in vivo*. This is consistent with reports demonstrating that histone acetylation upstream of promoters is limited to those with divergent transcription^33–35^. Second, comparison of the distributions of Epl1 and H4K12ac in actively transcribing cells, showed markedly different patterns of enrichment. For genes with unidirectional promoters, so as to avoid the confounding effect of divergent transcription, we found Epl1 bound upstream of promoters and throughout gene bodies while H4K12ac was primarily enriched in 5’ regions (Figure 3A). Consistent with this, Epl1 occupancy poorly correlated over genome-wide nucleosome positions with H4 acetylation marks (Figures 3B and S4A). Similar results were observed following treatment of cells with TSA (Figures 3A, B and S4A), and thus the differences between HAT occupancy and acetylated histone levels was not due to HDACs reshaping acetylation patterns. Finally, a similar disconnect was observed between occupancy of the H3-specific HAT, Gcn5^36^, and histone H3 acetylation (Figures 3B and S4A), suggesting that HAT binding poorly predicts histone acetylation for both H3 and H4 HATs. We note however that this observation could not be generalized to other chromatin-modifying enzymes, as the occupancy of the H3K4 methyltransferase, Set1, positively correlated with H3K4me3 (Figure 3B and S4B). Thus, specifically for HATs, we observe poor correspondence between HAT localization and histone acetylation, which is consistent with previous suggestions that regulation of HAT activity, following chromatin binding, is a major determinant of histone acetylation genome-wide^21, 26, 37, 38^.

**Figure 3.**
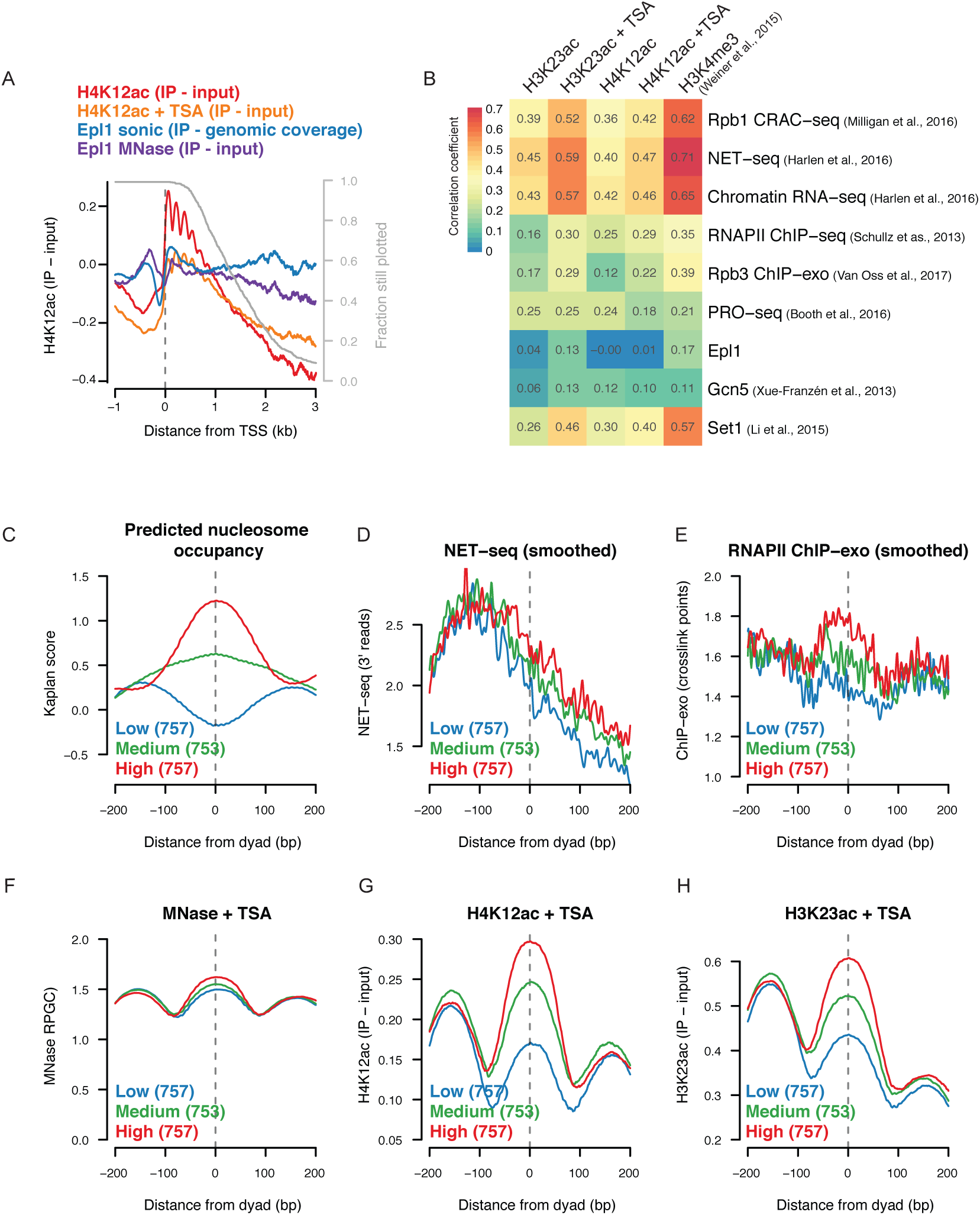
The activity of histone acetyltransferases is regulated post-recruitment. **A**. Average profiles of Epl1 (sonicated and MNase ChIP-seq) and H4K12ac (MNase ChIP-seq) at 709 unidirectional genes aligned by the TSS. Data from MNase treated extracts are shown as enrichment relative to nucleosome occupancy. **B**. Spearman correlation for nucleosome-normalized H3K23ac and H4K12ac (MNase ChIP-seq) with various measures of transcription across genome-wide nucleosome positions (Weiner et al., 2015). **C-H**. The average enrichment relative to +2, +3, and +4 NCP dyad positions for predicted nucleosome occupancy (**C**, Kaplan et al., 2009), NET-seq (**D**, Harlen et al., 2016), RNAPII ChIP-exo (**E**, Van Oss et al., 2017), nucleosome occupancy (**F**, MNase-seq), nucleosome-normalized H4K12ac (**G**) and H4K12ac (**H)** MNase ChIP-seq from TSA-treated cells. Nucleosome positions were filtered to remove those with high or low MNase-seq signal and “High” and “Low” were classified as those with a Kaplan score greater and 0.5 or less than -0.25 respectively.

High resolution mapping of engaged RNAPII by NET-seq, CRAC-seq, and chromatin bound RNA-seq, shows RNAPII accumulation at the 5’ ends of genes^33, 39, 40^, which is proposed to represent slow RNAPII passage through these regions. Similar 5’ accumulation is not observed with other high-resolution analyses of RNAPII presence, including ChIP-exo^41^, which simply measures RNAPII binding to DNA, or PRO-seq^42^, which measures elongation competent RNAPII. Interestingly, histone acetylation correlates strongly with NET-seq, CRAC-seq, and chromatin bound RNA-seq (Figure 3B), especially following TSA treatment, supporting the hypothesis that histones are acetylated by available HATs in regions with slow RNAPII passage. To test this, we asked whether nucleosomes predicted to impede RNAPII movement exhibit increased histone acetylation in vivo. We scored the +2, +3, and +4 nucleosomes relative to the TSS based on their predicted ability to strongly or weakly form nucleosomes (Figure 3C)^43^. Upstream of the analyzed nucleosomes, RNAPII signals were similar, indicating equivalent levels of transcription initiation at these genes (Figure 3D and E). In contrast, nucleosome-favoring sequences had increased NET-seq and RNAPII ChIP-seq signal within the nucleosome (Figure 3D and E), indicative of impaired elongation and consistent with in vitro data suggesting that DNA sequence can determine the strength of the nucleosomal impediment to RNAPII passage^44^. Importantly, intrinsically stable nucleosomes were enriched for H3K23ac and H4K12ac relative to total nucleosomes, both under steady state and TSA-treated conditions (Figure 3G and H, S5). These data further demonstrate the tight link between RNAPII and histone acetylation and suggest that strongly positioned nucleosomes can regulate both RNAPII elongation and histone acetylation in vivo.

## Discussion

In this study, we found that the majority of histone acetylation is dependent on transcription and is targeted to nucleosomes at sites of RNAPII accumulation. Histone acetylation is proposed to facilitate RNAPII elongation by directly modulating histone-DNA contacts or by targeting chromatin remodelers to disrupt nucleosomes^45^. Thus, our research suggests that acetylation is a component of a feed forward loop that maintains expression of active genes.

To understand the mechanism for targeting acetylation we mapped occupancy of Epl1, a component of NuA4 and Piccolo, and found that transcription is required to target this protein to active genes. Similar results were observed with Tip60, the mammalian homolog of Esa1, the catalytic subunit of NuA4 and Piccolo^46^. NuA4 consists of 12 subunits, including multiple putative targeting modules^47^. The largest subunit, Tra1, is proposed to mediate the interaction of multiple HATs with transcription activators^7–9^. A second subunit, Eaf3, preferentially interacts with H3K36-methylated histones^48, 49^, a post-translational modification associated with transcription elongation^2^. Surprisingly however, disruption of NuA4 did not deplete Epl1 from the bodies of active genes suggesting that the HAT complex bound in this region is Piccolo. Piccolo is made up of Esa1, Epl1 and Yng2^28^, and although Yng2 contains a methylated H3K4 binding motif^31^, we show that H3K4 methylation on its own was insufficient to target histone acetylation. Thus, at this time, the mechanism for targeting HATs to transcribed genes is unclear. Tip60 recruitment is proposed to occur via R-loops^46^, which has not been reported before for NuA4. However, as Esa1 contains a nucleic acid binding domain^50, 51^, this may be a conserved mechanism of HAT recruitment to transcribed regions.

Although histone acetylation requires the interaction of HATs with chromatin, genome-wide HAT occupancy is a poor predicter of histone acetylation, indicative of additional levels of regulation. One possible mechanism is via the alteration of nucleosome structure during transcription. The histone tails impede RNAPII progression through the nucleosome and thus these peptides must be displaced from DNA during elongation^52^, which could make them a better substrate for acetylation. The slow passage of RNAPII through the 5’ regions of genes increases the opportunity for HATs to access the histone tails resulting in increased acetylation in these regions. In support of this mechanism, the H3 tail becomes ten-fold less accessible when assembled into nucleosomes *in vitro*^53^ and HAT activity is enhanced by disrupting interactions between DNA and the H3 tail^54^. Moreover, we show that nucleosome favoring sequences tend to accumulate both RNAPII and histone acetylation.

Another possible mechanism for regulation of HAT activity is that RNAPII, or an associated factor, regulates the histone acetyltransferase activity of HAT complexes. This may be an allosteric regulator, as has been observed for enhancer RNAs and CBP^21^, but altered subunit composition is another promising mechanism of regulation. Esa1 exhibits reduced nucleosome HAT activity when incorporated into NuA4 when compared to Piccolo^28^. Similarly, Gcn5 nucleosomal HAT activity is relatively inefficient when incorporated into SAGA than when present in the smaller ADA complex^55^. Since NuA4 and SAGA complexes target Esa1 and Gcn5 respectively to DNA-bound transcription activators, the reduced HAT activity of these complexes may explain why increased targeting doesn’t translate into increased histone acetylation. Similarly, NuA4 is targeted to nucleosomes marked with H3K36me3^32, 49^, which tend to be hypoacetylated. Thus, the targeting of the different HAT complexes to specific regions may dictate the extent to which the nucleosomes present are modified.

Recruitment of HATs to promoters by transcription activators underlies the model that histone acetylation primarily occurs upstream of transcription initiation. However, whether HAT recruitment aids transcription initiation through histone acetylation or through some other function has largely been unexplored. We found that increased NuA4 recruitment to promoters did not lead to increased histone acetylation, and generally observed that HAT association with chromatin in the absence of RNAPII, rarely led to efficient histone acetylation. This suggests that HATs recruited by sequence-specific activators alone is not sufficient for histone acetylation. This does not rule out the possibility that activator-targeted HATs do acetylate histones following transcription initiation and nucleosome disruption. However, decades of research point to an important role for NuA4, and other HATs, in initiation transcription, so if not histone acetylation what is the function of HATs in this process? While activator targeting alone does not lead to histone tail acetylation, acetylation of other targets may be important for transcription initiation. In addition to the histone tails, subunits of the RSC and SWI/SNF chromatin remodeling complexes are known to be acetylated, as are many other factors involved in transcription^14^, and these represent potential targets for non-histone acetylation. Supporting a non-histone tail acetyltransferase function for HATs in transcription initiation, *in vitro* transcription of some chromatin templates can be stimulated by recruitment of p300 even when chromatin is assembled using tail-less histones^56^. The transcription stimulation depends on acetyltransferase activity, and radiolabeled acetate incorporation shows acetylation of many non-histone proteins. CBP and p300 were recently shown to acetylate numerous proteins involved in transcriptional regulation at enhancers and promoters in mammalian cells^57^, meaning that transcription activation in response to p300 recruitment^58^ cannot be assumed to be mediated through histone acetylation. Compared to the histone tails the role of non-histone acetylation is a relatively unexplored avenue of research, and our results highlight its potential importance in transcription initiation.

## Acknowledgments

Support for this work was provided by grants to L.J.H. and M.C.L. from the Canadian Institutes of Health Research (PJT-162253) and Natural Sciences and Engineering Research Council (RGPIN-2018-04907). B.J.E.M. was supported by a fellowship from the Natural Sciences and Engineering Research Council. We are grateful to Steven Hahn, Michael Kobor and Hiroshi Kimura for providing plasmids and antibodies. We are also grateful to Jerry Workman and Swami Venkatesh for useful discussions.

## Author Contributions

Conceptualization, L.J.H. and B.J.E.M.; Methodology, Software, Formal Analysis, and Visualization, B.J.E.M.; Investigation, B.J.E.M., J.B., A.K., K.N., J.L.; Writing – Original draft, B.J.E.M.; Writing – Reviewing & Editing, B.J.E.M., L.J.H., J.B., and M.C.L.; Supervision and Funding Acquisition, L.J.H. and M.C.L.

## Methods

### Cell Culture

FUCCI reporter mESCs^59^ were grown in standard feeder-free conditions in complete mESC media: Dulbecco Modified Eagle’s Medium (DMEM) high glucose, 15% fetal bovine serum (HyClone Laboratories), 20 mM HEPES, 1 mM L-glutamine, 100 U/ml penicillin-streptomycin,1 mM nonessential amino acids, ∼10-50 ng/ml of recombinant LIF, 1 mM sodium pyruvate and 0.1 mM β-mercaptoethanol on 0.2% type A gelatinized tissue culture plates.

### Yeast strains and growth

All strains used in this study were isogenic to S288C and are listed in table S3. Yeast culture and genetic manipulations were performed using standard protocols. Genomic deletions were verified by PCR analysis and whole cell extracts were generated as previously described^60^.

### Drug treatments

Yeast drug treatments were performed in YPD media at the following concentrations: 1,10 phenanthroline monohydrate (400 μg/mL in ethanol; Sigma 161-0158), thiolutin (10 μg/mL in DMSO; Santa Cruz SC-200387), 1-naphthalene acetic acid (1 mM in 85% ethanol; Sigma N0640), doxycycline (20 μg/ml in 50% ethanol; Sigma D9891), 1-Naphthyl PP1 (5 μM in DMSO; Sigma CAS 221243-82-9), trichostatin A (25 μM in DMSO), α factor (10 μM in 100 mM sodium acetate, pH=5.2, Sigma custom synthesis of peptide: WHWLQLKPGQPMY). mESCs were treated with Actinomycin D at 25 μg/mL (in DMSO; Sigma CAS 50-76-0).

### Immunoblot analysis

Whole cell lysates or cellular fractions were analyzed by SDS-PAGE using the antibodies listed in the key resource table. Blots were scanned and fluorescent signal quantified using the Licor Odyssey scanner.

### ChIP-seq

Yeast cells, grown to mid-log, were arrested in G1 by 3 hour treatment with 10 μM alpha factor. Cell synchronization was verified by cell “shmooing,” as seen under the microscope. For transcription inhibition, cells were treated with 400 μg/mL 1,10 phenanthroline monohydrate or 25 μM TSA for 15 minutes. Cells were crosslinked in 1% formaldehyde for 15 minutes and quenched with addition of liquid glycine to 125 mM for a further 15 minutes. Cells were lysed by bead beating, and cell lysate was spun down at 15,000g for 30 minutes.

For sonicated ChIP-seq, the pellet was resuspended in lysis buffer (50 mM HEPES, pH 7.5, 140 mM NaCl, 0.5 mM EDTA, 1% triton X-100, 0.1% sodium deoxycholate) and sonicated (Biorupter, Diagenode) to produce an average fragment size of 250 bp. The lysate was spun down at 9,000g for 10 minutes, and the supernatant was precleared by rotating with Protein G Dynabeads for 1 hour at 4 °C. Twenty percent of the lysate was reserved for input, and the remaining was split into two and incubated with α-HA antibodies overnight at 4°C.

For MNase ChIP-seq, the pellet was resuspended in MNase digestion buffer (0.5 mM spermidine, 1 mM β-ME, 0.075% NP-40, 50 mM NaCl, 10 mM Tris pH 7.4, 5 mM MgCl_2_, 1 mM CaCl_2_). Samples were incubated with 100 units of MNase for 10 minutes at 37°C. Lysates were clarified by centrifugation at 9000g for 10 minutes. To extract insoluble chromatin, pellets were re-suspended in 200 μL of lysis buffer with 0.2% SDS and sonicated in a Diagenode Bioruptor at medium output for 30 seconds on and 30 seconds off for four cycles, before centrifugation at 9000g for 10 minutes. The second supernatant was pooled with the first, and the buffer composition of the lysate was adjusted to that of the original lysis buffer (50 mM HEPES pH 7.5, 140 mM NaCl, 1 mM EDTA, 2% Triton X-100, 0.2% Na-deoxycholate, 1X Roche protease inhibitor cocktail, 1 mM PMSF). The supernatant was precleared by rotating with Protein G Dynabeads for 1 hour at 4°C, ten percent of the lysate was reserved for input, and immunoprecipitations were performed using α-HA, α-H3K23ac, α-H4K12ac, or α-H4K8ac, antibodies.

Antibody immunoprecipitations were isolated by adding magnetic Protein G Dynabeads and rotating at 4°C for 1 hour, and 5 minute washes were performed twice with lysis buffer, twice with high salt buffer (50 mM HEPES pH 7.5, 640 mM NaCl, 1 mM EDTA, 2% Triton X-100, 0.2% Na-deoxycholate), twice with LiCl wash buffer (10 mM Tris-HCl pH 8.0, 250 mM LiCl, 0.6% NP-40, 0.5% Na-deoxycholate, 1 mM EDTA), and once with TE. Synthetic spike-in DNA was added to eluates, to aid in quantification.

Following proteinase K digestion, DNA was purified by phenol, chloroform, isoamyl alcohol (PCI) extraction and RNase A treated.

### ChIP-seq Library Preparation

Libraries for paired-end sequencing were constructed essentially as described previously^61^, using a custom procedure for paired-end sequencing. Briefly, 2–10 ng of ChIP material was end-repaired and A-tailed before being ligated to TruSeq PE adaptors. The adaptor-ligated material was subject to 8 to 11 rounds of PCR amplification, and an aliquot of each library was run on an Agilent Tape Station to check the size distribution and molarity of the PCR products. Equimolar amounts of indexed, amplified libraries were pooled, and fragments in the 200–600 bp size range were selected on an agarose gel. An aliquot (1 μL) of the library pool was run on an Agilent Tape Station to confirm proper size selection. In between each reaction, the material was purified using NucleoMag solid phase reversible immobilization paramagnetic (SPRI) beads.

### Analysis of ChIP-seq data

Adapter sequences were removed from paired end fastq files using cutadapt (version 1.83– http://cutadapt.readthedocs.io/en/stable/), before aligning to the saccer3 genome using BWA (version 0.7.15-r1140)^62^. Coverage tracks represent reads per genome coverage, calculated using deeptools version 3.02^63^. For IP over input tracks, a value of 1 coverage per million fragments was added to both to avoid division by 0, and the resultant track was smoothed with a 10 bp moving average window and log_2_ transformed. Replicates were pooled for subsequent analysis, and figures were generated in R.

Similar to other groups^25, 26^, ChIP-seq datasets comparing across RNAPII perturbations were normalized to silent regions. The genome was divided into 250 bp bins, bins outside the interquartile range for coverage in the input were discarded, the 100 regions with the lowest Rpb3 signal were defined as silent regions, and these silent regions were used to normalized ChIP-seq datasets for cross-condition comparisons (Table S2). We also added synthetic DNA spike-ins to our ChIP eluates and inputs (Table S3), but this approach to normalization did not work well for all samples, possibly due to low coverage of the spike-ins in some samples.

For transcribed nucleosomes classified by Rpb3 change upon 1,10-pt treatment (Figure S1F), genome-wide nucleosome positions^64^ with Rpb3 signal greater than the median were classified as transcribed. Nucleosomes where Rpb3 changed by less than 10% were classified as “Rpb3 stable”, while those decreasing by at least 3x were classified as “Rpb3 lost”. Boxplots represent the 1^st^ to 3^rd^ quartiles, with whiskers extending to 1.5 times the interquartile range or to the extreme of the data. Notches are equal to +/-1/58 IQR /sqrt(n), giving an approximation of the 95% confidence interval for the difference in 2 medians.

### Publicly available datasets

Publicly available datasets used in this paper are listed in Table S4. SRA files were downloaded and converted to fastq using fastq-dump from the SRA toolkit (version 2.8.2-1). Fastq files were mapped to the saccer3 genome using BWA version 0.7.15-r1140^62^. For single end sequencing experiments, reads were extended to the reported fragment length, and reads per genome coverage were calculated using deeptools version 3.02^63^. For histone PTM ChIP-seq data^64^, IP over input tracks were calculated as per the “Analysis of ChIP-seq” section. For Rpb3 ChIP-exo experiments^41^, the crosslinking point was taken as the first nucleotide sequenced from the 1^st^ read of the read pair. The Java Genomics Toolkit (https://github.com/timpalpant/java-genomics-toolkit) was used to generate wig coverage tracks (ngs.BaseAlignCounts) of crosslinking points, pool replicates (wigmath.Average), and apply a Gaussian smoothing curve using a standard deviation of 3 bp (wigmath.GaussianSmooth). For enrichments over nucleosome positions, within 50 bp of each annotated nucleosome dyad^64^ the number of fragment midpoints per million reads were calculated. For histone PTMs these were normalized by matched MNase inputs or sonicated bulk histone IPs. Nucleosome positions with no coverage in the MNase inputs were removed.

### Defining genome annoations

Transcription start and end sites were downloaded from the supplemental files of Chereji et al., 2018^65^. Transcribed genes were defined as genes in the top 3 quartiles for RNAPII ChIP-seq reads in 5’ regions (TSS to +500 bp). Further as we observed enrichment of Epl1 and our untagged HA ChIP-seq IPs at tRNAs and centromeres respectively, genes within 500 bp of tRNA genes or centromeres were removed, giving 3862 TSSs for metaplots. For analysis of *epl1.485* data, genes on chrXII were removed as well, as this chromosome appeared to be unstable in this mutant (1.5x coverage) giving 3520 TSSs. Unidirectional promoters were defined as having low NET-seq^40^ signal from -100 to -750 bp from the TSS, giving 709 TSSs. For heatmaps, which were sorted by the change in RNAPII upon transcription inhibition, genes were considered irrespective of RNAPII enrichment, giving 5133 TSSs. For +2, +3, and +4 NCPs, called nucleosome positions from chemical mapping of dyads^66^ were assigned a nucleosome position relative to the TSS^65^. Nucleosomes positions were then filtered by MNase-seq coverage, keeping nucleosomes in the middle two quartiles. Non-transcribed nucleosomes were removed from analysis, using a cutoff of an average of 0.25 NET-seq^40^ reads over the nucleosome. Then low, medium, and high predicted nucleosome occupancy scores were selected as quantile groups 0-20, 40-60, and 80-100 percent respectively.

### Generating heatmaps and metaplots

To make metaplots and heatmaps, deeptools was used to generate matrices of the various data tracks centred on TSSs or NCPs. These matrices were loaded into R (version 3.5.2) and metaplots and heatmaps were generated using baseR and ComplexHeatmap version 1.20^67^ respectively. For 2D heatmaps, matrices were generated using Bedtools version 2.27.1^68^ and Bedops version 2.4.30^69^, and heatmaps were plotted in R using pheatmap (version 1.0.12).

### Data availability

The ChIP-seq data have been deposited in the Gene Expression Omnibus (https://www.ncbi.nlm.nih.gov/geo/) under accessions GSE110286 and GSE110287.

---

**Table S1:**
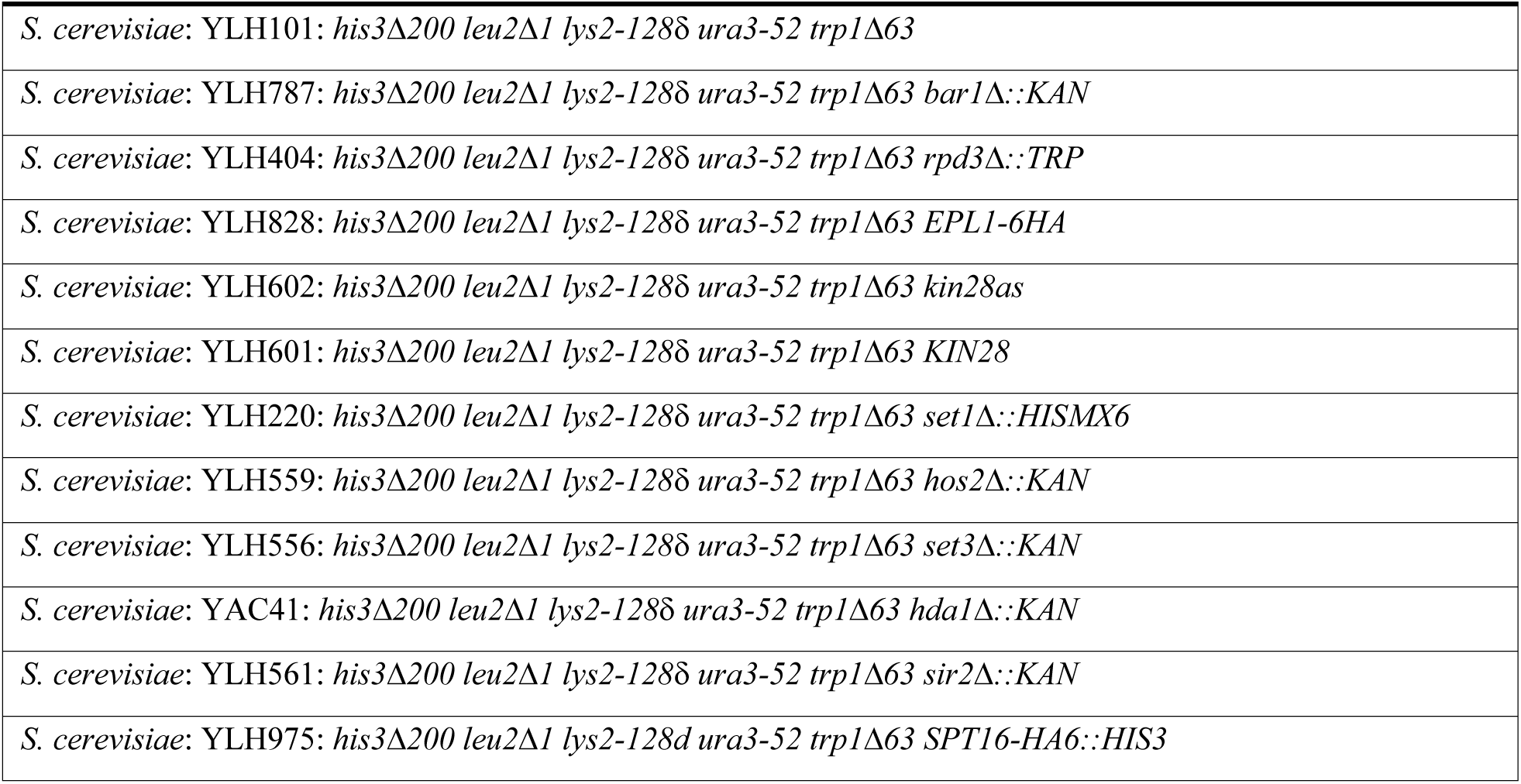
Yeast Strains.

**Table S2:** Antibodies.

|  |  |  |
| --- | --- | --- |
| H3K4me3 | Abcam | ab1012, lot #1276040 |
| H3K9ac | Custom antibody (Kimura et al., 2008) | CMA305 |
| H3K14ac | Custom antibody (GeneScript)<sup>61</sup> | Affinity-purified rabbit polyclonal antibody |
| H3K18ac | Abcam | ab1191 |
| H3K23ac | Active Motif | 39131, lot # 1008001 |
| H4K5ac | Millipore | 07-327, lot # 2524676 |
| H4K8ac | Abcam | ab45166, clone # EP1002Y |
| H4K12ac | Active Motif | 39165, lot # 1008001 |
| H4K16ac | Millipore | 07-329, lot # 2506422 |
| H3 | Custom antibody (GeneScript)<sup>61</sup> | Affinity-purified rabbit polyclonal antibody, Raised against sch3 peptide (CKDILARRLRGERS) |
| H4 (C-terminal region) | Abcam | ab31830, myeloma: Sp2/0-Ag14 |
| IgG | Millipore | PP64, lot # LV1602263 |
| HA | Roche | 12CA5, lot # 11849700 |
| 3e8 (ser5p) | Millipore | 04-1572, lot # 2585825 |

**Table S3:**
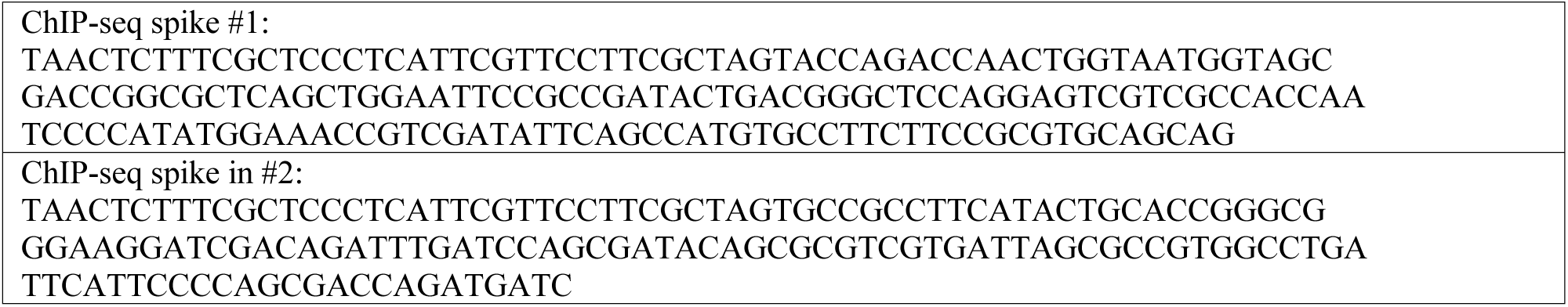
ChIP-seq spike-ins (IDT gblocks)

**Table S4:**
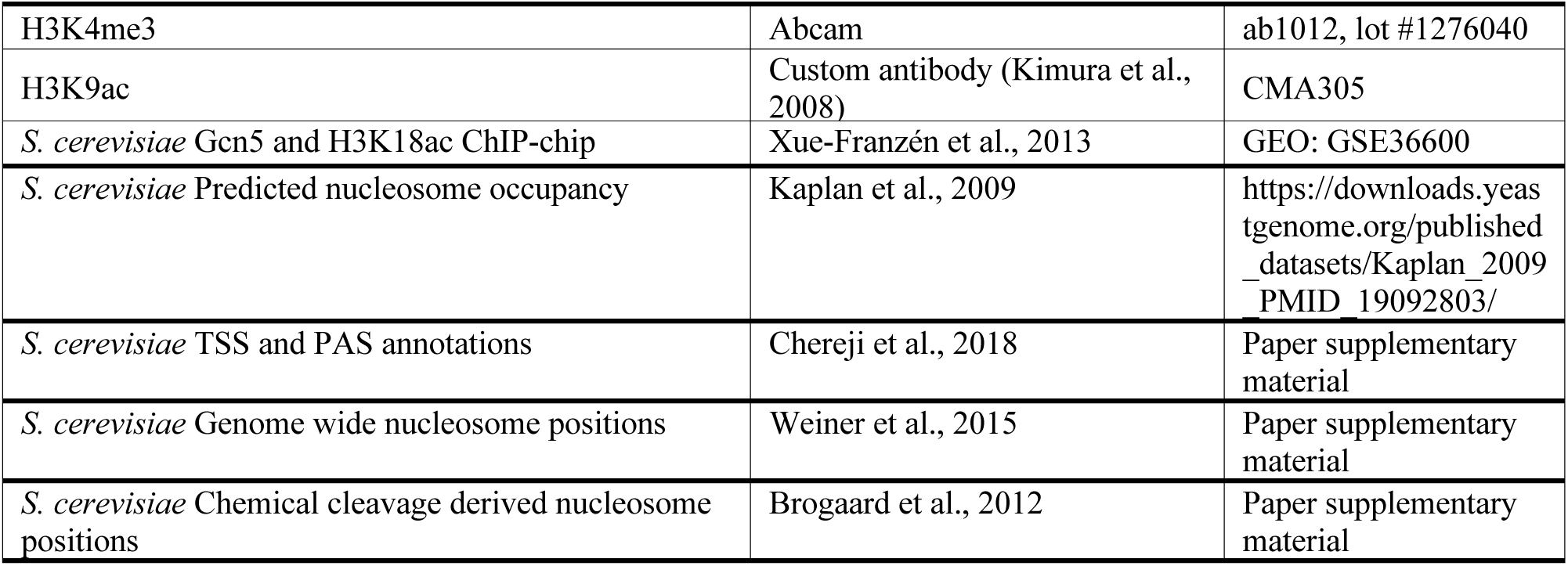
Published datasets used.

## Supplemental Figure Legends

**Figure S1:**
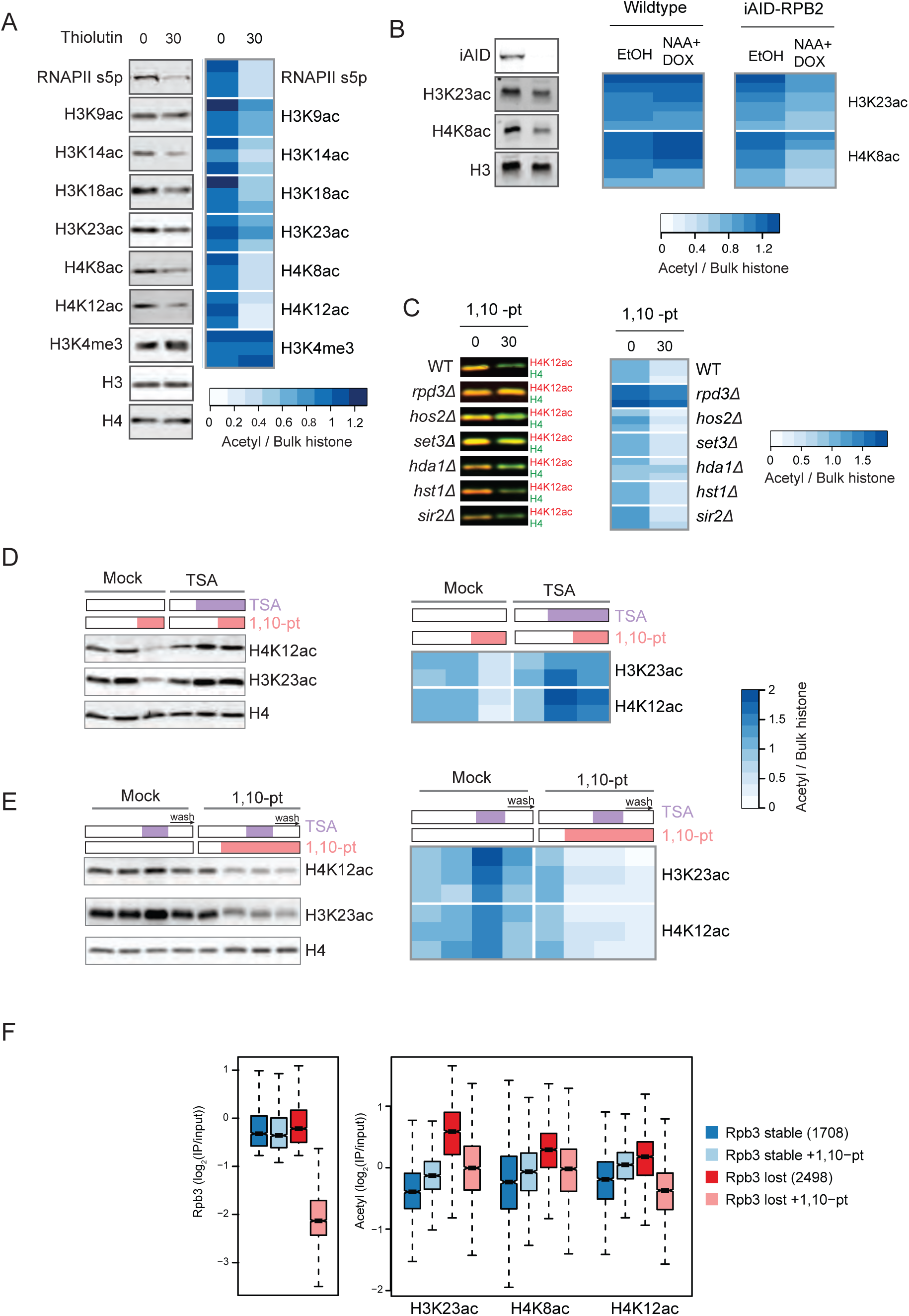
Characterization of histone deacetylation following transcription inhibition. **A**. Whole cell extracts from the *S. cerevisiae* cells before and after treatment with thiolutin were subjected to immunoblot analysis with the indicated antibodies. Representative blots (left panel) and quantification of three independent replicates (right panel) are shown. For quantification, histone PTM signals were normalized to total histone H4 levels and expressed as a heatmap with the scale below. **B**. Whole cell extracts from the indicated strains after treatment with ethanol or auxin and doxycycline were subjected to immunoblot analysis with the indicated antibodies. Representative blots (left panel) and quantification of six independent replicates (right panel) are shown. For quantification, histone PTM signals were normalized to total histone H3 levels and expressed as a heatmap with the scale below. **C**. Wildtype and deacetylase mutant strains were treated with 1,10-pt for 30 minutes and bulk H4K12ac was assessed by immunoblotting. Representative blots are shown (left panel) as well as quantification of three replicates (right panels). For quantification, histone PTM signals were normalized to total histone H4 levels. **D**. Cells were pretreated with the HDAC inhibitor trichostatin A (TSA) or DMSO for 15 minutes before a 30-minute treatment with 1,10-pt. Whole cell extracts were subjected to immunoblot analysis and representative blots (left panel) and quantification of two replicates (right panel) are shown. For quantification, histone PTM signals were normalized to total histone H4 levels. **E**. Cells were pretreated with 1,10-pt or ethanol for 15 minutes, followed by a 15-minute treatment with TSA, and washed into fresh media containing 1,10-pt or ethanol without TSA for 15 minutes. Whole cell extracts were subjected to immunoblot analysis. Shown are representative blots (left panel) and quantification of three replicates (right). For quantification, histone PTM signals were normalized to total histone H4 levels. **F.** RNAPII (Rpb3 ChIP-seq) and nucleosome-normalized histone acetylation (MNase ChIP-seq) enrichments at transcribed nucleosomes with stable RNAPII and those that lost RNAPII upon treatment with 1,10-pt.

**Figure S2:**
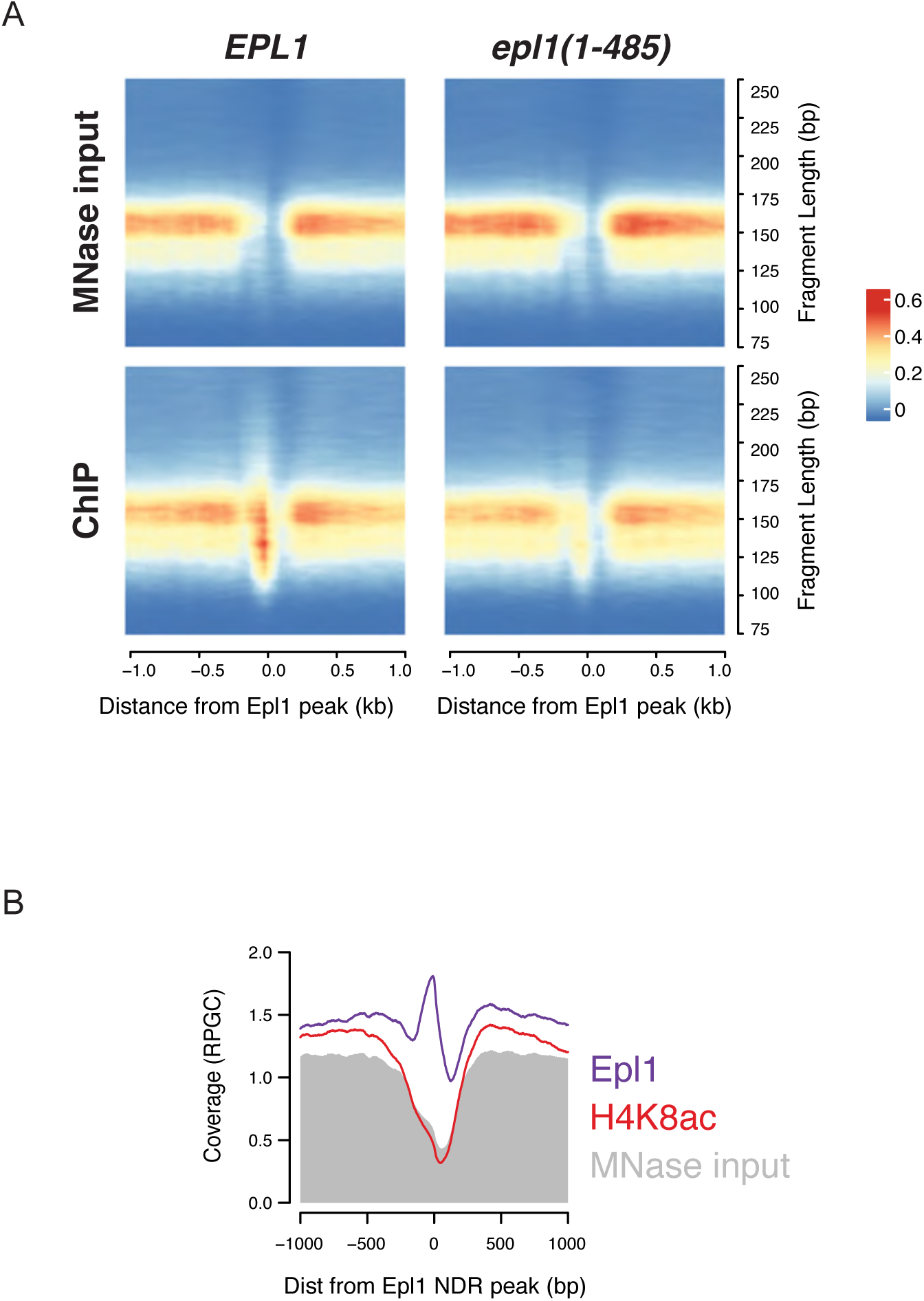
Promoter-localized Epl1 peaks are dependent on NuA4 and not associated with acetylation. **A.** Two-dimensional occupancy plots of relative sequence fragment abundance, sequence fragment length, and sequence fragment position from input and Epl1 ChIP from MNase-digested chromatin, relative to the center of 562 Epl1 peaks. Plot was generated using plot2DO^70^ run with standard settings. The relative sequence read abundance is indicated as a heatmap, the sequence fragment length is plotted on the y-axis, and the position of sequence reads relative to the peak center plotted on the x-axis. **B.** Average profile of input, Epl1 ChIP and H4K5ac ChIP from MNase-digested chromatin relative to the center of 562 regions showing strong Epl1 peaks.

**Figure S3:**
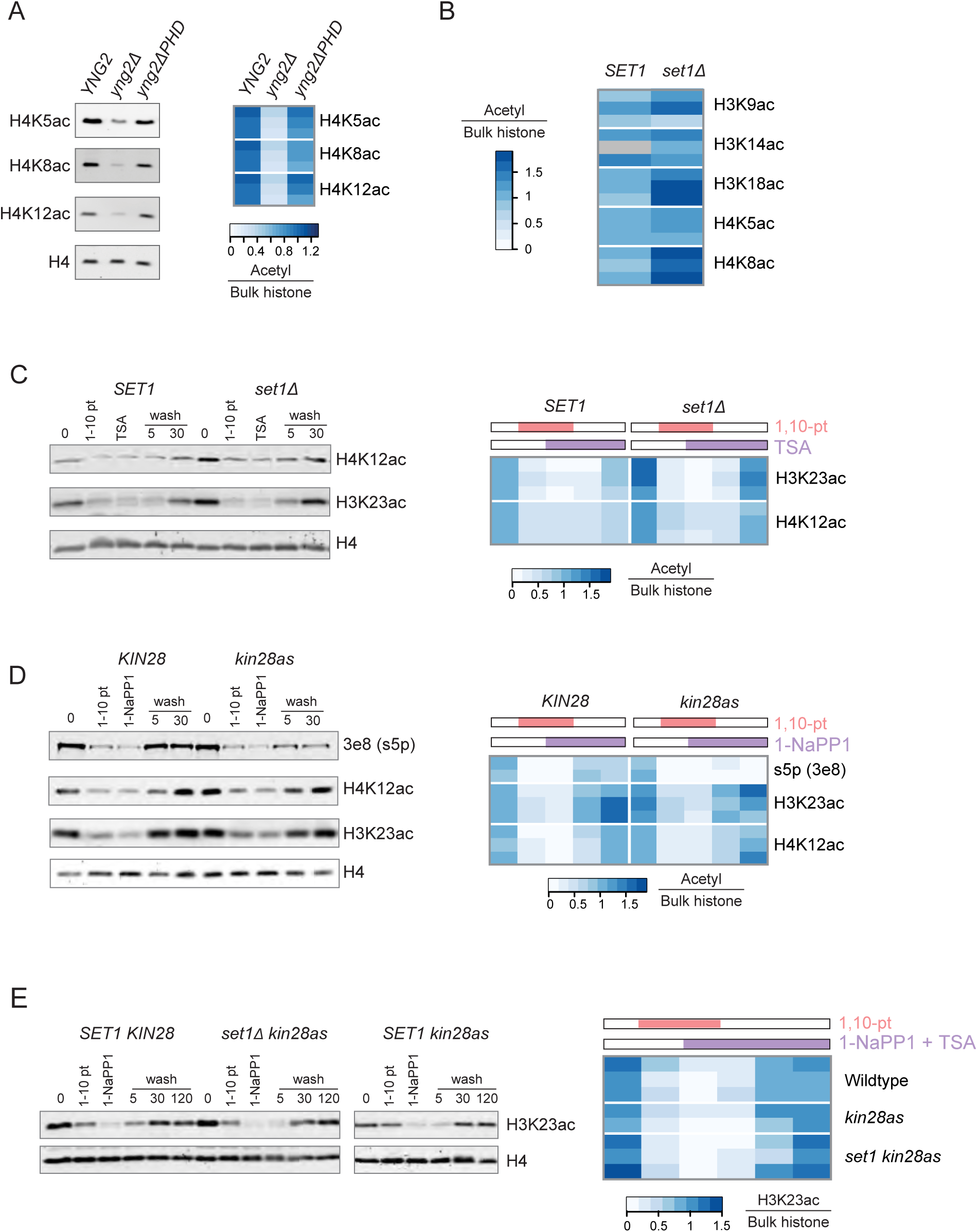
Known pathways for targeting histone acetylation are not required for bulk H3K23 or H4K12 acetylation. **A**. Bulk H4 acetylation levels in wild type, *yng2*Δ, and *yng2*Δ*PHD* cells. Shown are representative blots (left panel) and quantification of three independent replicates by heatmap (right panel). **B**. Bulk histone acetylation levels in wildtype and *set1*Δ strains were assessed by immunoblotting and quantification of three replicates is shown. A gray box indicates a missing value for H3K14ac. **C**. Immunoblot analysis of wild type and *set1*Δ cells treated with 1,10-pt for 30 minutes, followed by TSA treatment for an additional 30 minutes, before being washed into fresh media containing TSA. Samples were collected 5- and 30-minutes post-wash. Shown are representative blots (left panel) and quantification of three replicates by heatmap (right panel). **D**. Immunoblot analysis of wild type and *kin28as* strains treated with 1,10-pt for 30 minutes, followed by 1-NaPP1 for an additional 20 minutes, before being washed into fresh media containing only 1-NaPP1. Samples were collected 5- and 30- minutes post-wash. Shown are a representative blot (left panel) and quantification of three replicates by heatmap (right panel). **E**. Immunoblot analysis of wild type, *kin28as,* and *kin28as set1*Δ cells treated with 1,10-pt for 30 minutes, followed by TSA treatment for an additional 30 minutes, before being washed into fresh media containing TSA. Samples were collected 5, 30, and 120 minutes post-wash. Shown are representative blots (left panel) and quantification of three independent replicates by heatmap (right panel). For quantification, H3K23ac of all immunoblots, signals were normalized to total histone H4 levels.

**Figure S4:**
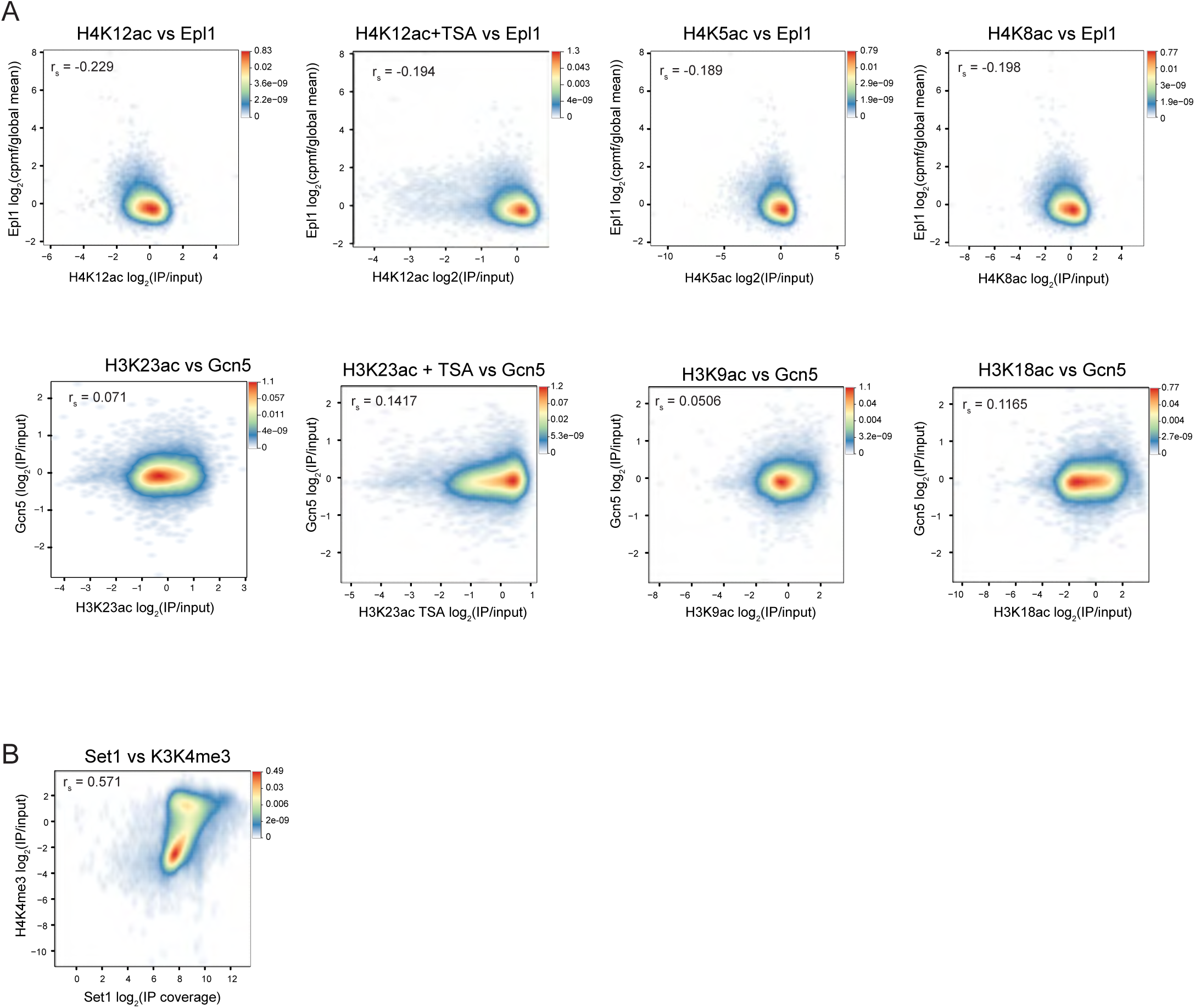
HAT occupancy poorly correlates with histone acetylation. **A.** Smoothed scatter plots across genome-wide nucleosome positions^64^ for Epl1 (ChIP-seq) and Gcn5 (ChIP-chip^71^) versus nucleosome-normalized H3K9ac (ChIP-chip^71^), H3K18ac (ChIP-chip^71^), H3K23ac, H4K5ac, H4K8ac, and H4K12ac with and without TSA treatment as indicated. **B.** Smoothed scatter plots across genome wide nucleosome positions for Set1 (ChIP-seq^72^) versus nucleosome-normalized H3K4me3 (MNase ChIP-seq).

**Figure S5:**
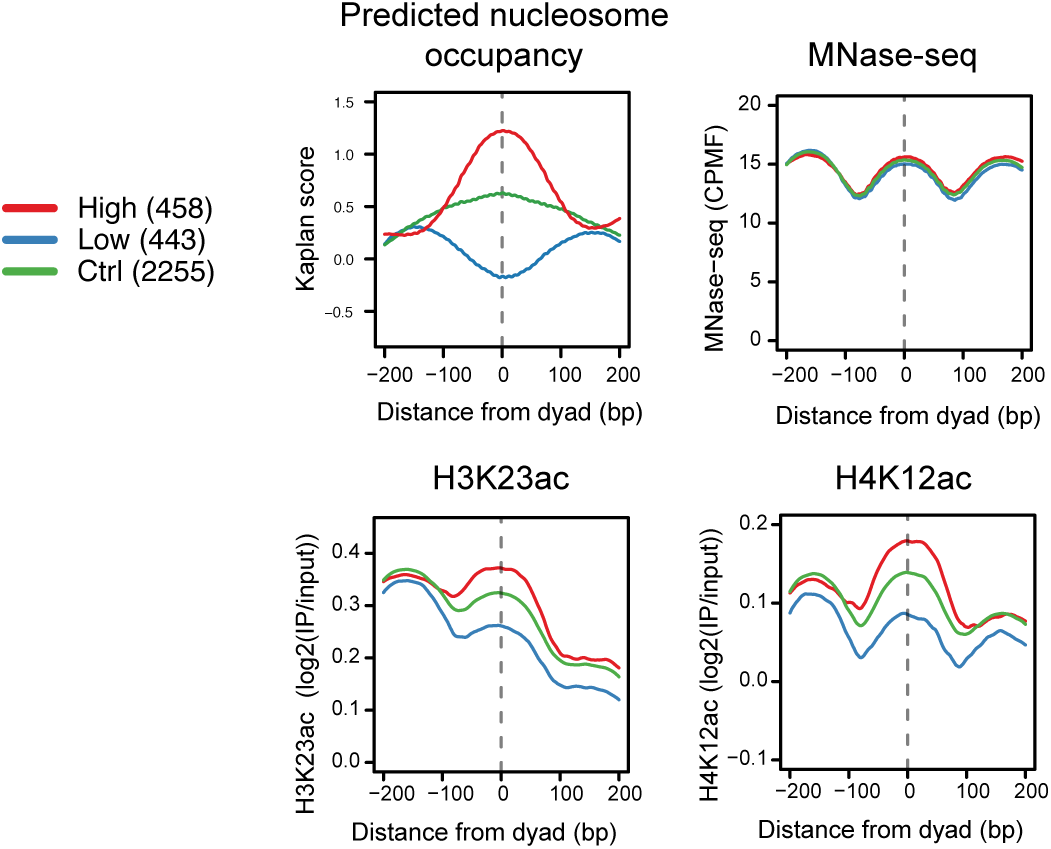
Histone acetylation is strongly linked with stalled and backtracked RNAPII. The average enrichment relative to +2, +3, and +4 NCP dyad positions for predicted nucleosome occupancy^43^, nucleosome occupancy (MNase-seq), nucleosome-normalized H4K12ac and H3K23ac (MNase ChIP-seq) from non-TSA-treated cells. Nucleosome positions were filtered to remove those with high or low MNase-seq signal and “High” and “Low” were classified as those with a Kaplan score greater and 0.5 or less than -0.25 respectively.

